# Development of EST-SSR annotated database in olive (*Olea europaea*)

**DOI:** 10.1101/026294

**Authors:** Sami S. Adawy, Morad M. Mokhtar, Alsamman M. Alsamman, Mahmoud M. Sakr

## Abstract

Olive tree (*Olea europaea* L.) is one of the most important oil producing crops in the world and the genetic identification of several genotypes by using molecular markers is the first step in its breeding programs. A set of 1,801 well-informative EST-SSR primers targeting specific Olive genes included in different biological processes and pathways were generated using 11,215 Olive EST sequences acquired from the NCBI database. Our bioinformatics analytical procedure showed that 8295 SSR motifs were detected which belonged to different motif types with occurrences of 77.6%, 11.84%, 8.62%, 0.84%, 0.77% and 0.29% for Mononucleotide, trinucleotide, dinucleotide, hexanucleotide, pentanucleotide and tetranucleotide respectively. The appearance of the AAG/CTT repeat was highly represented in trinucleotide and the representation of AG/CT was high in dinucleotide repeats. Results obtained from functional annotation of olives EST sequences targeted with our primers set indicated that 78.5% of these sequences having homology with known proteins, while 4.2% was homologous to hypothetical, predicted, unnamed or uncharacterized proteins and the 17.3% sequences did not possess homology with any known proteins. Our EST-SSR primer set cover a total of 92 biological pathways such as carbohydrate metabolism pathway, energy metabolism& carbon fixation in photosynthetic organism pathway including 11 pathways associated with lipid metabolism. A twenty five randomly selected primers were applied to 9 Egyptian cultivated olive accessions to test its amplification and polymorphism detection efficacy. All tested primers were successfully amplified and only 10 exhibited detectable polymorphism.

## Introduction

Olive tree (*Olea europaea* L.) is one of the most superannuated and important long lived fruit species in Mediterranean (Zohary et al. 2012), it is a diploid species (2n = 2x = 46) with a genome size ranging between 2.90 pg/2C and 3.07 pg/2C, with 1C = 1,400 - 1,500 Mbp (Loureiro et al. 2007). *Olea europaea* is one of the first domesticated crops from *Oleaceae* family for oil production and the second most important oil fruit cultivated crop worldwide (Baldoni et al. 2009).Olive is a dependable source of edible oil and food for several thousands of years (Newton et al. 2006; Ben-Ayed et al. 2014; Calzada et al. 2015). The large number of accessions cultivated in olive producing countries make the olive germplasm preservation and management a major problem as far as olive breeders are concerned (Awan et al. 2011).

The development for early selection strategies in olive breeding programs is a main goal at present (Atienza et al. 2014) and in this view, using molecular markers techniques for the identification and characterizing of several genotypes is the first step in modern olive breeding programs (Bracci et al. 2011) and choosing a co-dominant, reliable and well amplified marker type is very crucial to start this process in order to significantly minimize the quantity of breeding starting materials and promotes the selection of desirable genotypes, which posses desired genes in its homozygous state (Sivolap 2013).

Reflecting its increasing rate of mutation, micro-satellites repeats shows a highly level of length polymorphism (Sahu et al. 2012) with a high evolution rates and a possible impact on the modification genes they are associated with. Not to mention that the typical role of mutation is to add or subtract repeat units which are both reversible and frequent, making SSR influence on genes regulation depending on the repeats number and provide a source of qualitative and quantitative variations (Kashi and King 2006).

These features granted SSR derived techniques its high heterozygosity (Powell et al. 1996; Adam-Blondon et al. 2004; Luro et al. 2008) and the ability to differentiate between different accessions with distinct agronomical advantages, despite synonymous problems in many plant species (Díaz-Losada et al. 2012; Trujillo et al. 2013; Vantini et al. 2015). This arise the need of developing new derived SSR markers with a PCR primers rich resources more linked to desired genic regions in different plant species, mean while the improvement and increasing of DNA sequencing technologies aid the increasing and sequencing of expressed genes was used to construct a large collection of EST libraries isolated from different tissues of various organism under distinct environmental conditions and through different development stages (Ozgenturk et al., 2010).

Recent studies reported the using EST libraries as a reliable resource for SSR derived markers taking in advance the availability of EST sequences in public databases and bioinformatics tools which detected SSR repeats and developed a PCR-based EST-SSR markers could reveal a high polymorphism in genic regions related to important agronomic traits (Gupta and Varshney 2000; KAUR et al. 2015). EST-SSRs markers reported in several plant species, such as *Musa* (Mbanjo et al. 2012), Finger Millet (Naga et al. 2012), *Jatropha Curcas* (Wen et al. 2010), Pineapple (Wöhrmann and Weising 2011), Citrus (Liu et al. 2013), Watermelon (Verma and Arya 2008), Sugarcane (Pinto et al. 2004), and bread wheat (Varshney et al. 2002).

In olive this technique could develop new functional markers with a flexibility to be used in marker-assisted selection in breeding programs and a useful tool for genes discovery, gene mapping, and gene-gene interaction, functional and comparative studies.

Sequence public databases contain a large number of EST sequences derived from different olive cultivars under a variety of environmental conditions, stand as useful resources for developing gene based markers. The aim of this study was to use bioinformatics analytical procedures to detect SSRs in Olive’s ESTs, compare the frequency and distribution of different repeat types in genic sequences, develop new genic EST-SSR markers suited for Olive genome, determine the localization of these primers targeted ESTs in different pathways and offer these primers in an informative illustration style to simplify the searching for trait - related markers in Olive breeding programs.

## Materials and Methods

A total of 11,215 *Olea europaea* ESTs sequences were acquired from NCBIEST database, these ESTs were isolated under distinct environmental conditions and through different developmental stages (http://www.ncbi.nlm.nih.gov).

SSRs identification was performed using the PERL script MISA (MIcroSAtellite identification tool; http://pgrc.ipk-gatersleben.de/misa/) and the criteria to determine SSR repeats were: mononucleotide (mono-) ≥ 10, dinucleotide (di-) ≥ 6, trinucleotide (tri-), tetranucleotide (tetra-), pentanucleotide (penta-), and hexanucleotide (hexa-) ≥ 5, and the number of maximum bases interrupting two SSRs to produce a compound microsatellite is 100 bp.

The flanking regions of SSR motifs were used to design SSR PCR-based primers using primer3_core (Untergasser et al. 2012). The parameters used: optimum length of primer was 20 nucleotides, optimum annealing temperature (Tm) of 58°C, expected to amplify products size of 100-500 bp and optimum G/C content of 50 %.

### Validation of designed primers

Twenty five PCR EST-SSR primers were randomly selected to validate its amplification efficacy, these EST-SSR primers were synthesized and applied on nine Olive cultivars adapted to the Egyptian environment (Maraki, Tofahi, Koratina, Pekoal, Manzanillo, Dolici, OjaziShami, Kronaki and Calamata).

Total genomic DNA was extracted from olive leaves using the Plant Genomic DNA Kit (Qiagen). PCR reaction content and PCR program cycles were summarized in **(File S1).**

### Olive ESTs GO enrichment analysis

Only Olive EST sequences contain detectable SSR motifs and has generated valid primers through previous mentioned criteria were used in GO enrichment analysis by using Blast2GO pipeline tool (Conesa et al. 2005) to assign gene ontology terms to EST products. BlastX search against the non-redundant (nr) NCBI database was used to analyze selected EST sequences with an Expect value (E-value) ≥1.0E-3 and the maximum hits for every gene was 20 hits. In the mapping and annotation steps of GO analysis, the default evidence codes weights (default=5) and Cutt-Off value score (default=55), respectively were used. The annotation step with GO-weight of 5 was given to map children terms of all EST sequences have hits.

## Results and Discussion

### Distribution of various repeat types of olive

Our result referred to 4,088 of *Olea europaea* EST sequences 36.45% out of 11,215 contains detectable SSR motifs matching our criteria, these ESTs contain 8,295 various SSR motifs. The gap between sequences contains simple repeats and repeat occurrence was due to the possibility that one SSR could contain more than one motif **(Table 1)**.

**Table 1.** Summary of SSR repeats identified on *Olea europaea* EST sequences.

| Searching item | Numbers |
| --- | --- |
| Total number of sequences examined: | 11215 |
| Total size of examined sequences (bp): | 6566149 |
| Total number of identified SSRs: | 8295 |
| Number of SSR containing sequences: | 4088 |
| Number of sequences containing more than one SSR: | 1910 |
| Number of SSRs present in compound formation: | 2447 |

Our investigation of different SSR repeats types showed that the highest appearance percentage of mono- repeats were 77.64%, followed by tri-11.84%, di-8.62%, hexa-0.84%, penta-0.77% and tetra-0.29% **(Figure 1).**The higher abundant of tri-in coding regions were consistent with previous studies in eukaryotic genomes (Jia et al. 2007; Rajendrakumar et al. 2008).

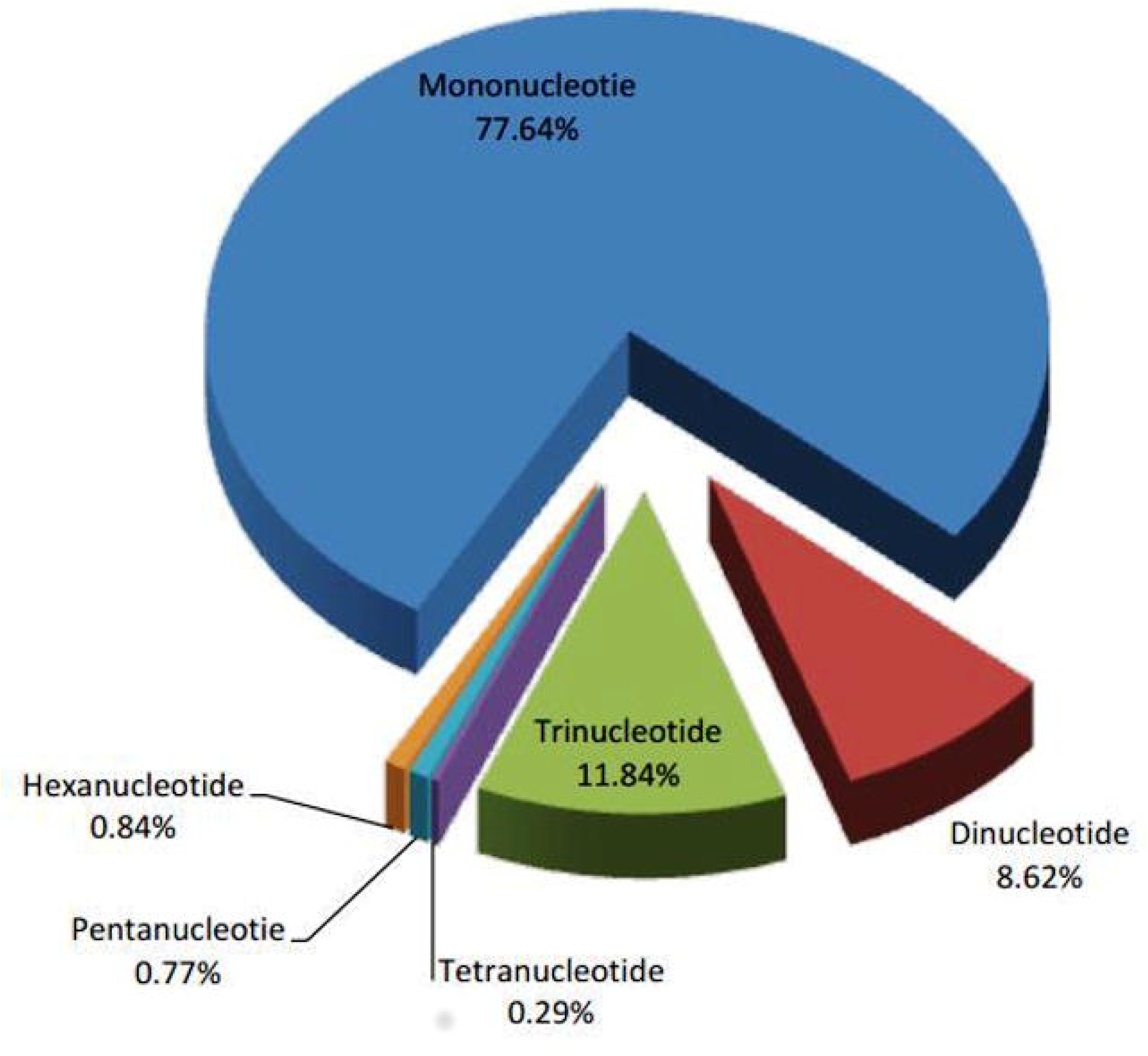

The mono- motifs A/T were 88.8% higher than G/C motifs (11.2%), and these results were proportionate with SSR analysis of chloroplast on *Olea* species (Filiz and Koc 2012) and with SSR analysis of major cereal organelle genome (Rajendrakumar et al. 2008). In di- motifs, GA represented 55% of the di- motifs in olive EST sequences, this agree with previous studies suggested that GA are the most abundant repeats type in foxtail millet (Jia et al. 2007), barley, maize, rice, sorghum and wheat (Kantety et al. 2002). AG/CT and GA/TC motifs were the most frequent respectively, while CG repeats were the lowest frequencies, this case was reported in microsatellites distribution for *Brassicaceae*, *Solanaceae* and *Poaceae* (Maia et al. 2009). The motifs Type of di- could represented in multiple codons depending on the open reading frame (ORF) regions which will be translated into different amino acids, for instance AG/CT motifs could represents AGA, GAG, CUC and UCU codons in mRNA, in this case it will be translated into the amino acids Glu, Arg, Leu and Ala respectively, therefore Ala and Leu will be presented in proteins at higher frequencies, hence the higher incidence of GA, CT motifs in the EST sequences (Lewin and Dover 1994). This could be one of the reasons suggested to explain the highly representation GA, CT motifs appearance in EST collections (Cho et al. 2000). di- repeats that located in coding regions are more sensitive to any change, such as substitutions, additions or deletions, as it causes a frame shifts which could give alternative amino acids (Metzgar et al. 2000). Regarding tri-, the TCT and TTC motifs were the most common repeats in olive EST **(Table 2)**, on the other hand AAG/CTT motifs were the most common in other studies focused on SSR types occurred in the chloroplast of *Olea* species (Filiz and Koc 2012), despite the fact that, CCG or AAC were the most common tri- repeats types in other crops such as barley, maize, rice, sorghum and wheat (Kantety et al. 2002).

**Table 2.**
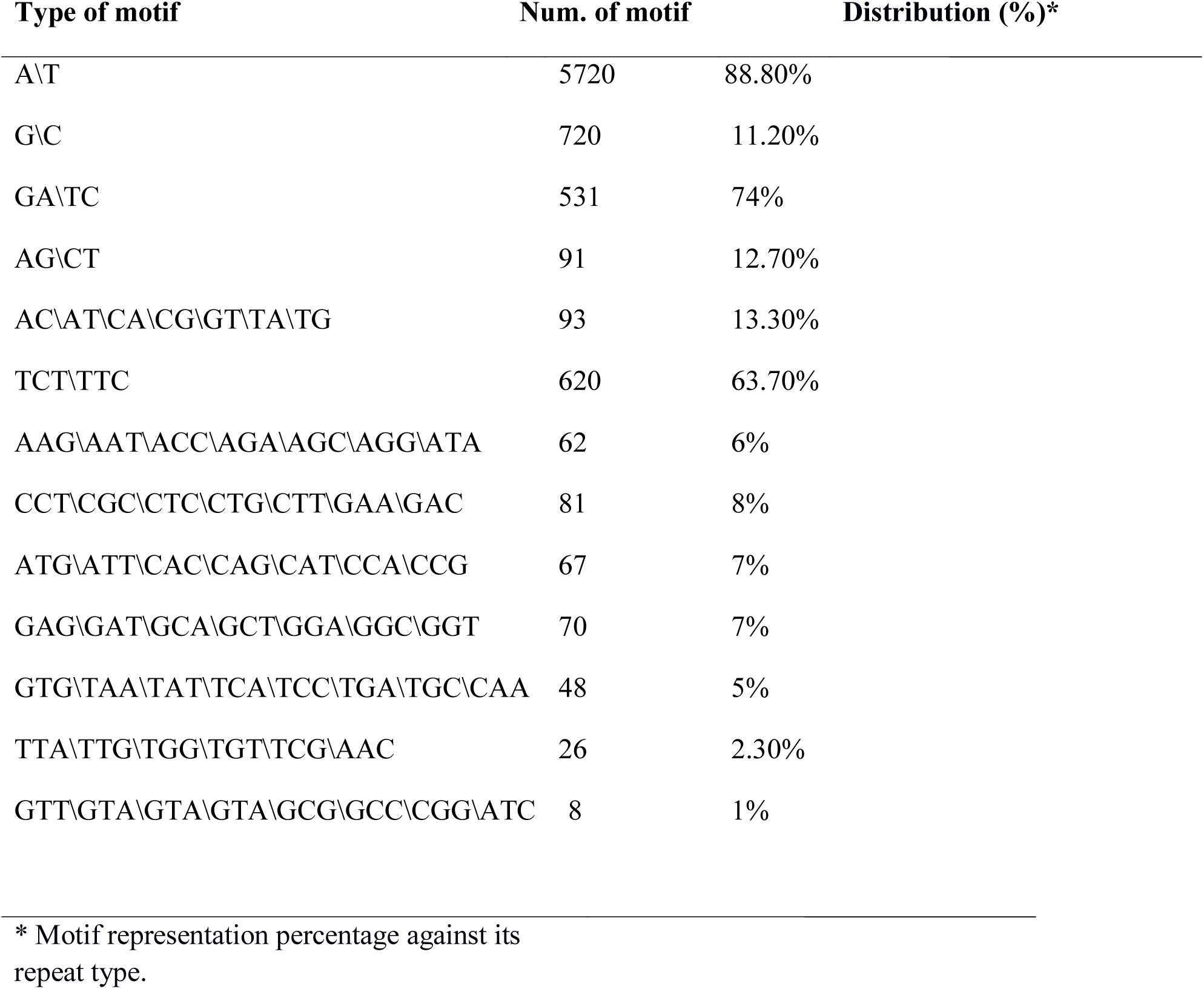
Distribution of different repeat types on *Olea europaea* EST sequences.

| Type of motif | Num. of motif | Distribution (%)* |
| --- | --- | --- |
| A\T | 5720 | 88.80% |
| G\C | 720 | 11.20% |
| GA\TC | 531 | 74% |
| AG\CT | 91 | 12.70% |
| AC\AT\CA\CG\GT\TA\TG | 93 | 13.30% |
| TCT\TTC | 620 | 63.70% |
| AAG\AAT\ACC\AGA\AGC\AGG\ATA | 62 | 6% |
| CCT\CGC\CTC\CTG\CTT\GAA\GAC | 81 | 8% |
| ATG\ATT\CAC\CAG\CAT\CCA\CCG | 67 | 7% |
| GAG\GAT\GCA\GCT\GGA\GGC\GGT | 70 | 7% |
| GTG\TAA\TAT\TCA\TCC\TGA\TGC\CAA | 48 | 5% |
| TTA\TTG\TGG\TGT\TCG\AAC | 26 | 2.30% |
| GTT\GTA\GTA\GTA\GCG\GCC\CGG\ATC | 8 | 1% |

\* Motif representation percentage against its repeat type.

Our results revealed that tetra-motifs AATC, CTTT are the most common; however the most common in *Olea* species SSRs chloroplast were AAAG, CTTT (Filiz and Koc 2012). Penta-AAAAT and hexa-GAAAAA were the most common motifs in our result, while AATCC was the most common on penta- in *Olea* species chloroplast and hexa- was not found in this organelle (Filiz and Koc 2012).

### EST-SSR PCR-based primer design

In this study, we used 4,088 EST sequences to design and select one of the most suitable PCR primer pairs. Only 1,801 EST sequences which contain detectable SSR motifs generated suitable primer pairs. The other ESTs 2287 sequences neither contain enough flanking regions to design a specific primer, or the generated primers didn’t match our criteria which we managed by primer3_core tool (Untergasser et al. 2012). The designed primers were referred as Oe-ESSR_xxxx, where Oe-ESSR is an abbreviating for *Olea europaea* EST-SSRs and xxxx are referring to the index of EST-SSR primers (start with 1 and end with 1801).

### Gene ontology enrichment analysis for Olive EST-SSR sequences

All EST sequences which have generated an EST-SSR primer pairs by our mentioned criteria were annotated with Blast2go pipeline tool. In the BLAST step, out of the 1,801 EST sequences used, only 1413 have a homology with known proteins, while hypothetical, predicted, unnamed or uncharacterized proteins were 75 and only 313 sequences did not possess homology with any known proteins. Most of these hits have Expected values ≥ 1.E-27 **(Figure 2-A)** and the homology degrees ranging from 40.5% to 100% **(Figure 2-B)**.

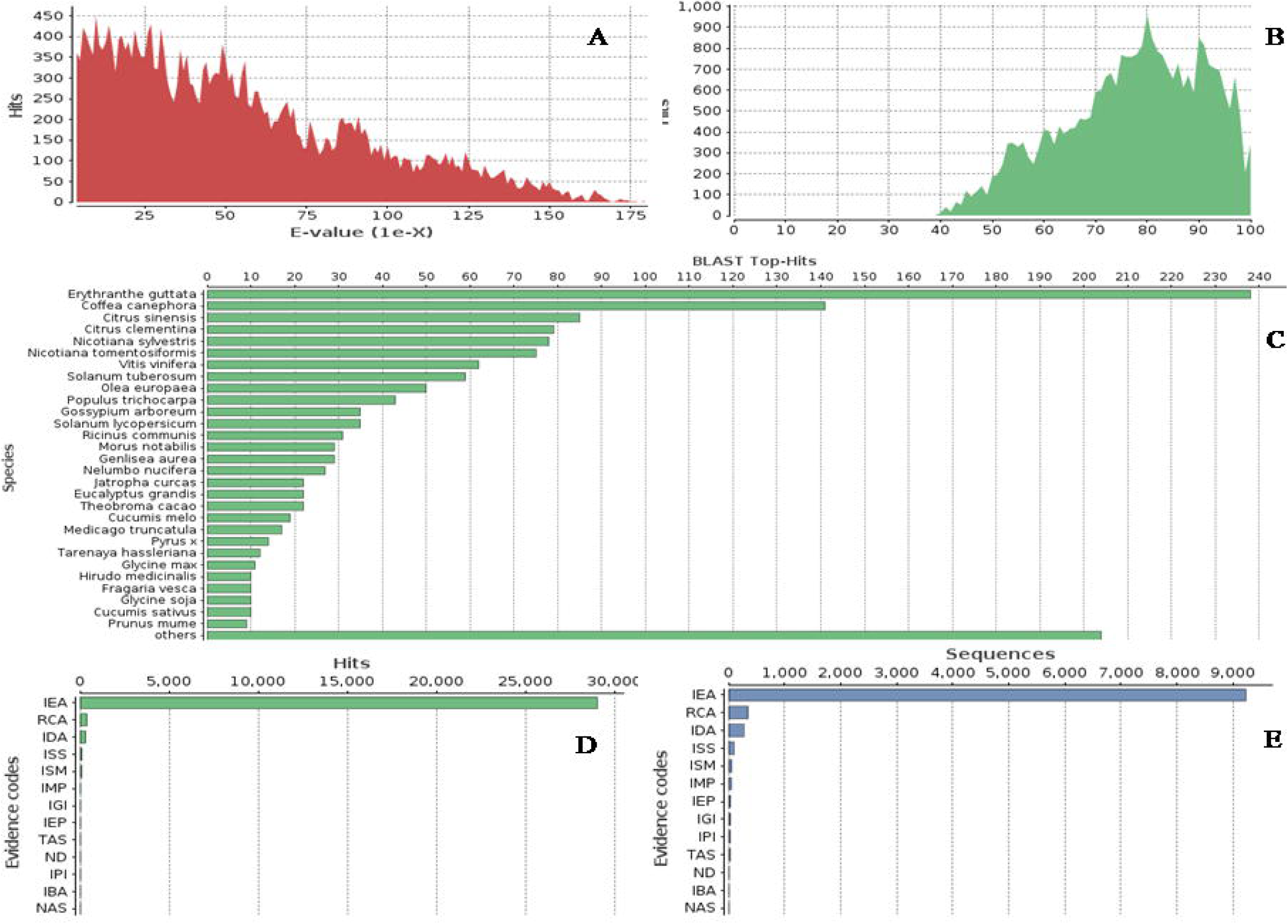

*Olea europea* came in the ninth place in top-hit species distribution, maybe due to that, only sequences revealed SSR and produced PCR primers were used or lower number of olive sequences in the NCBI database compared to other species with finished and published genomes **(Figure 2-C).**

In the GO terms mapping step, only 1264 sequences were mapped with a total of GO terms reaches 6432.The number of GO terms assigned to every EST sequence differs from one to 49 terms and most EST sequences were mapped to terms inferred from electronic annotation (IEA), which is higher in evidence code distribution for both blast hits and sequences **(Figure 2-D&2-E).**

In the annotation step, about 5090 GO terms were mapped to 1264 EST sequences, giving a GO mean level of 6.9 and revealing 256 sequences with known enzyme code (EC). The average length of sequences was 823 and sequences with length higher than 750 bp gain more annotation than other sequences. The other 537 EST sequences, which generated PCR primers and didn’t reveal any annotation could be used as a tool to discover genomic regions with unknown function.

The three major GO functional groups: molecular function (GO: 0003674), biological process (GO: 0008150) and cellular components (GO: 0008370) revealed subgroups with related biological functions. Out of 5090 GO terms revealed in our result, about 1348 are linked to molecular function, 1244 GO are related to cellular components and 2498 GO terms associated with biological processes **(Figure 3).**

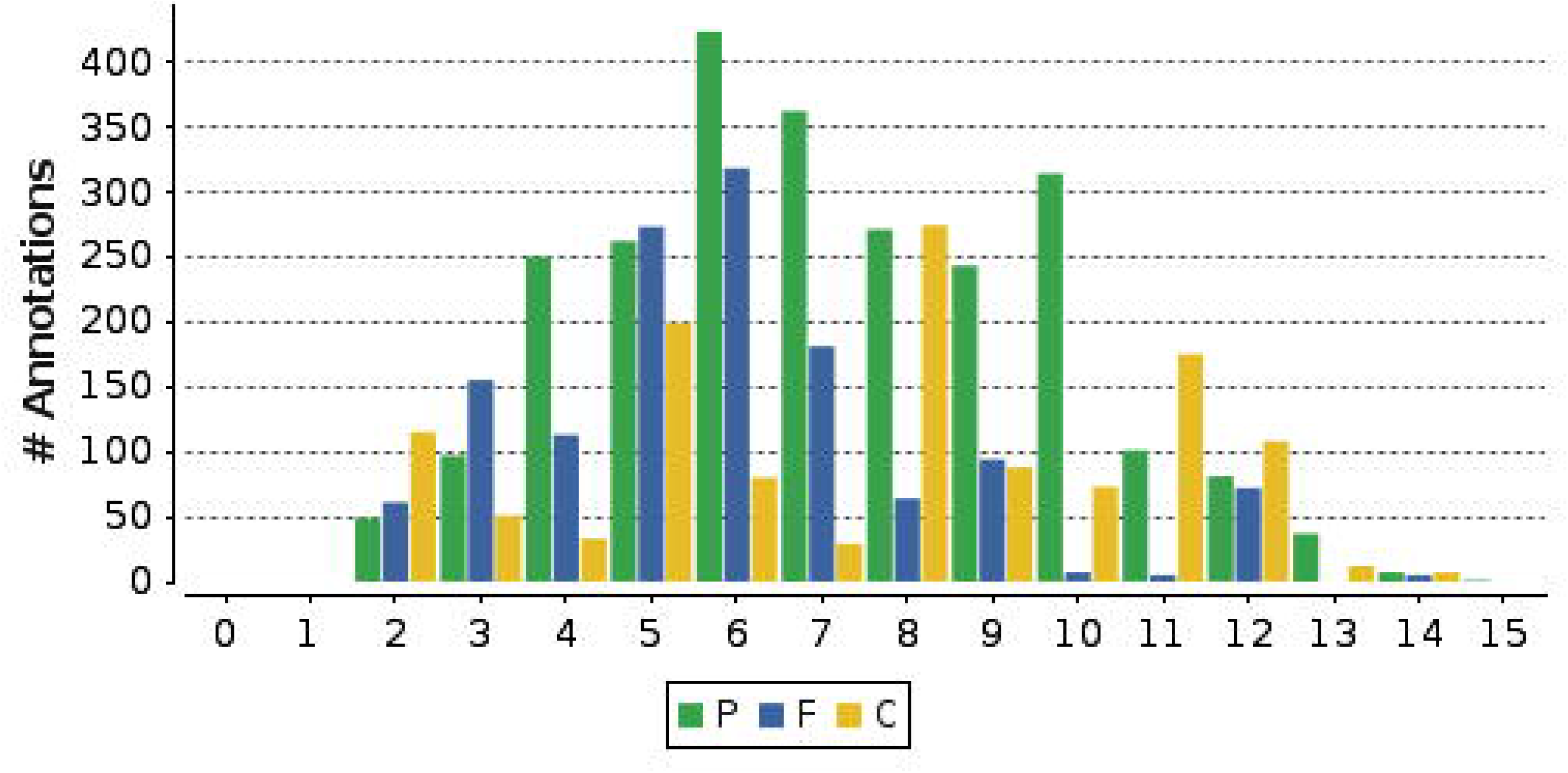

In the biological processes about 22% of the total EST-SSR sequences with PCR-based primers are associated with genes involving in cellular processes (GO:0009987) like cell communication, which its activation is reported under Olive environmental stresses and fruit development (Gucci et al. 2009; Hammami et al. 2011). Also metabolic processes (GO: 0008152) were covered with (21%) of ESTSSR primers, this processes involves beta-glucosidase, a gene that shaping the phenolic profile of virgin olive oil (Romero-Segura et al. 2012),

Other processes like single-organism process (GO:0044699) which includes genes that enhance the salt tolerance in some plant like *CIPKs* family (Hu et al. 2015),localization (GO:0051179), response to stimulus (GO:0050896) has gain 16%, 12%, 8% of ESTs, respectively, while signaling (GO:0023052), rhythmic processes (GO:0048511) and growth(GO:0040007) are covered with the lowest number of EST-SSR primers.

The molecular function category are covered with SSR primers targeting ESTs associated with catalytic activity (GO: 0003824) (37%), binding (GO: 0005488) (36%) including *SEUSS-LIKE* genes, which has been reported as transcriptional adaptors regulate the development of flower and embryo (Bao et al. 2010) and transporter activity (GO: 0005215) (16%) like aquaporin genes.

Cellular components category are assigned by cell (GO: 0005623) 42.7% primers targeting cell membrane genes and organelle (GO: 0043226) (13%) primers for organelle ESTs and macromolecule complex (GO: 0032991) (13%) **(Figure 4)**.

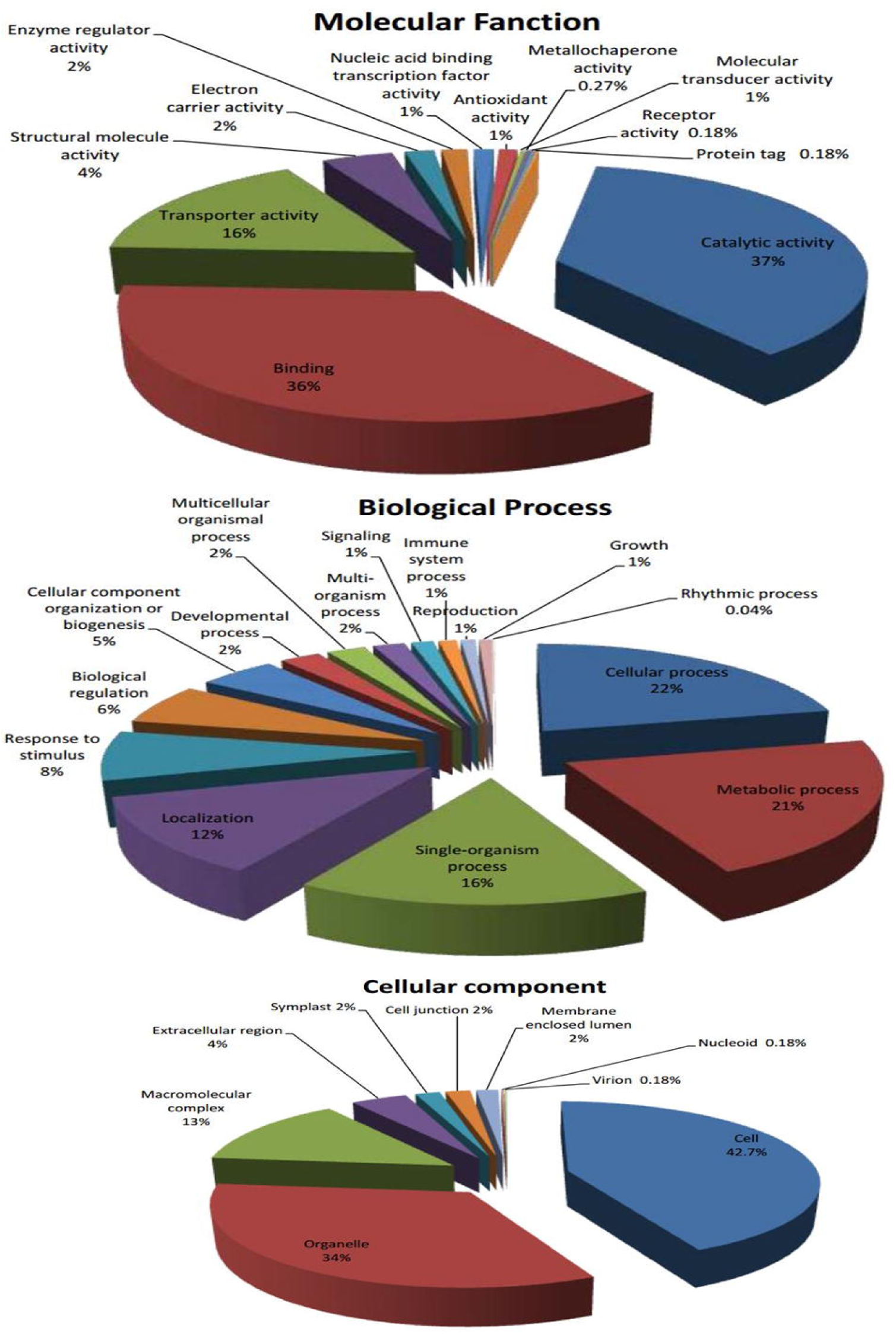

### Functional classification by KEGG pathway analyses

The KEGG pathway database is a useful tool for understanding genes biological functions and its molecular interactions (Li et al. 2012).To stand on the metabolic pathways that were covered by our EST-SSR primers set, we mapped all EST sequences that contains detectable SSR motifs and has generated valid primers to the KEGG reference pathways. The KEGG pathway analysis revealed that, our EST-SSR primers set covered a total of 92 different pathways and about 256 EST-SSR primers are associated with genes linked to 132 enzymes.

The major pathways were covered with EST-SSR primers using over than nine genes each. These pathways includes starch and sucrose metabolism which is related to depletion of stored carbohydrates (CHO) during the on-year (high yield) and suggested as a cause for alternate bearing in olives (Bustan et al. 2011).Another targeted metabolic pathway is gluconeogenesis which controls the manipulation of non-carbohydrate carbon substrates to glucose (Sung et al. 1988), also methionine metabolism which synthesized S-Adenosylmethionine as a donor of the methyl group in DNA methylation for gene expression regulation (Lu 2000). These ESTs has a significant match in the KEGG database **(Table S2)** and these results are visualized by using Circos software (Krzywinski et al. 2009) **(Figure 5).**

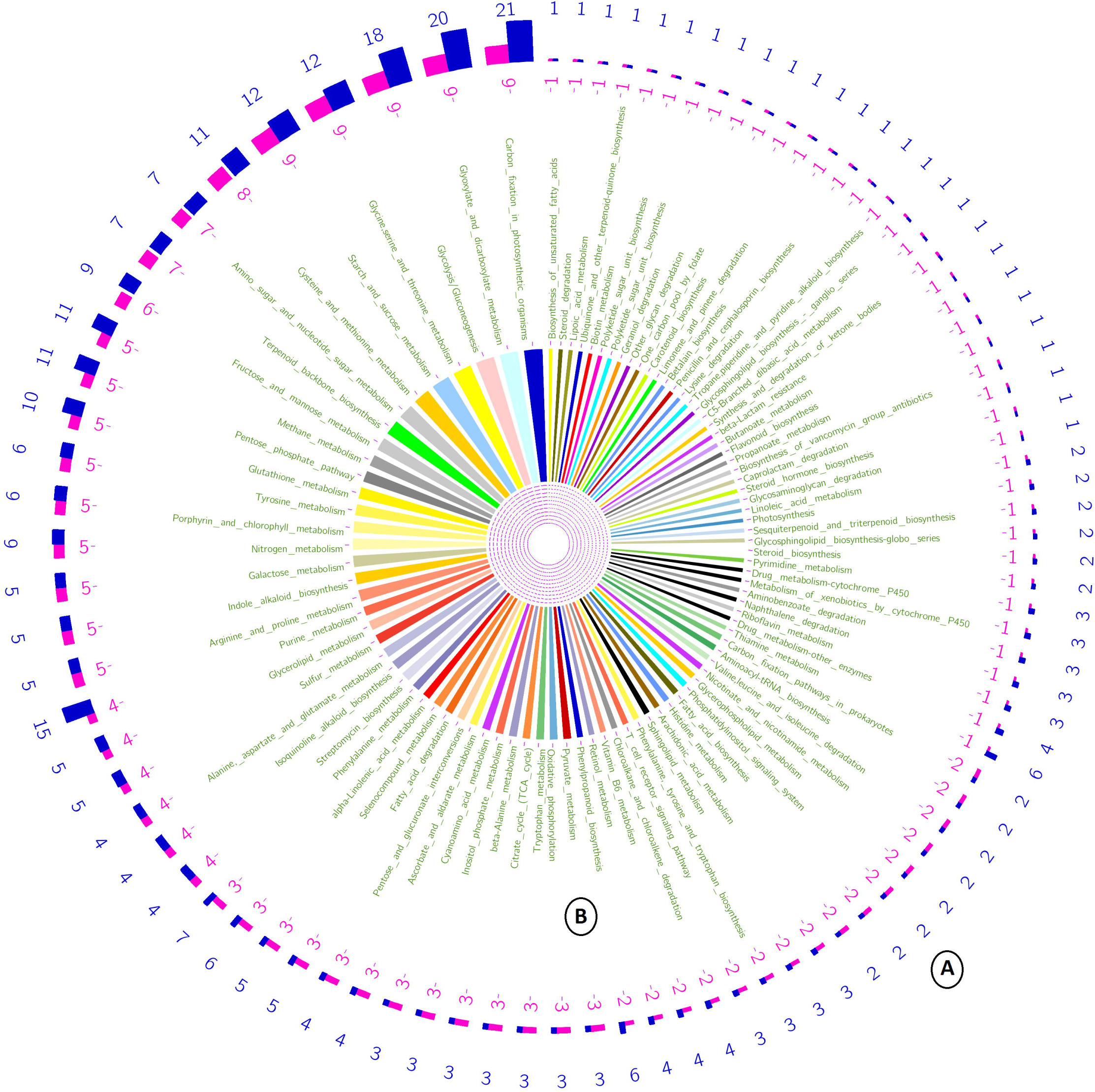

Breeding Olive trees for achieving a higher Olive oil quantity and quality is one of the most important goals for Olive breeding programs worldwide (El Riachy et al. 2012; Ozdemir et al. 2013).There is a high occurrence of EST-SSR primers in metabolic pathways for enzymes related oil contents indicates a good potential opportunity for using a marker type related to oil traits in olives. The primer-targeted ESTs were categorized by the metabolism it involves in, including lipid metabolism **(Table 3)**, carbohydrate metabolism, energy metabolism, amino acid metabolism, nucleotide metabolism and metabolism of cofactors and vitamins.

**Table 3.**
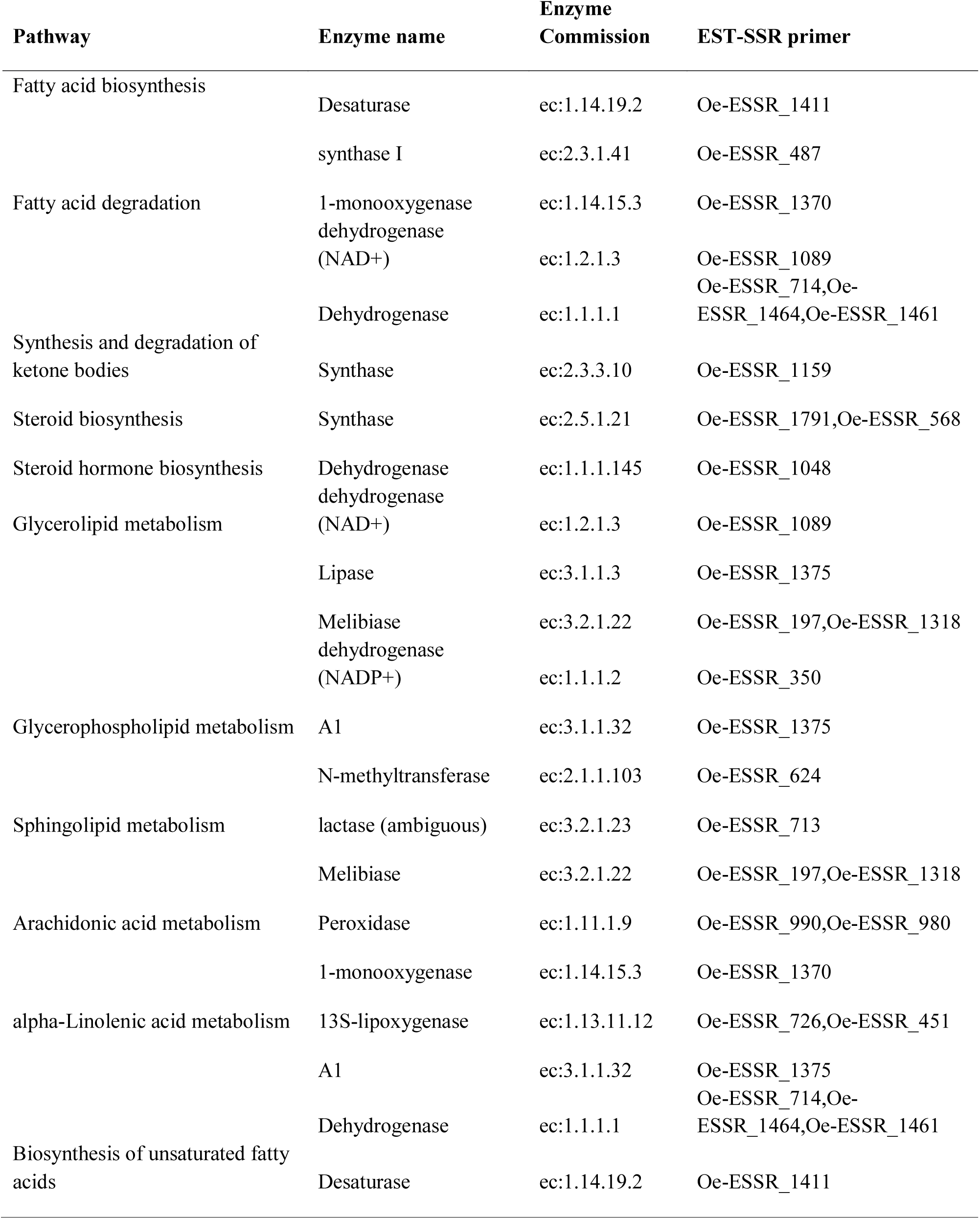
List of lipid metabolism pathways which have been assigned to EST-SSR sequences targeted with PCR primers.

In details, the mapping results can further investigated against the glycolysis/gluconeogenesis **(Figure 6)** and Fatty acid degradation pathways **(Figure 7)** as an example of carbohydrate metabolism and lipid metabolism respectively.

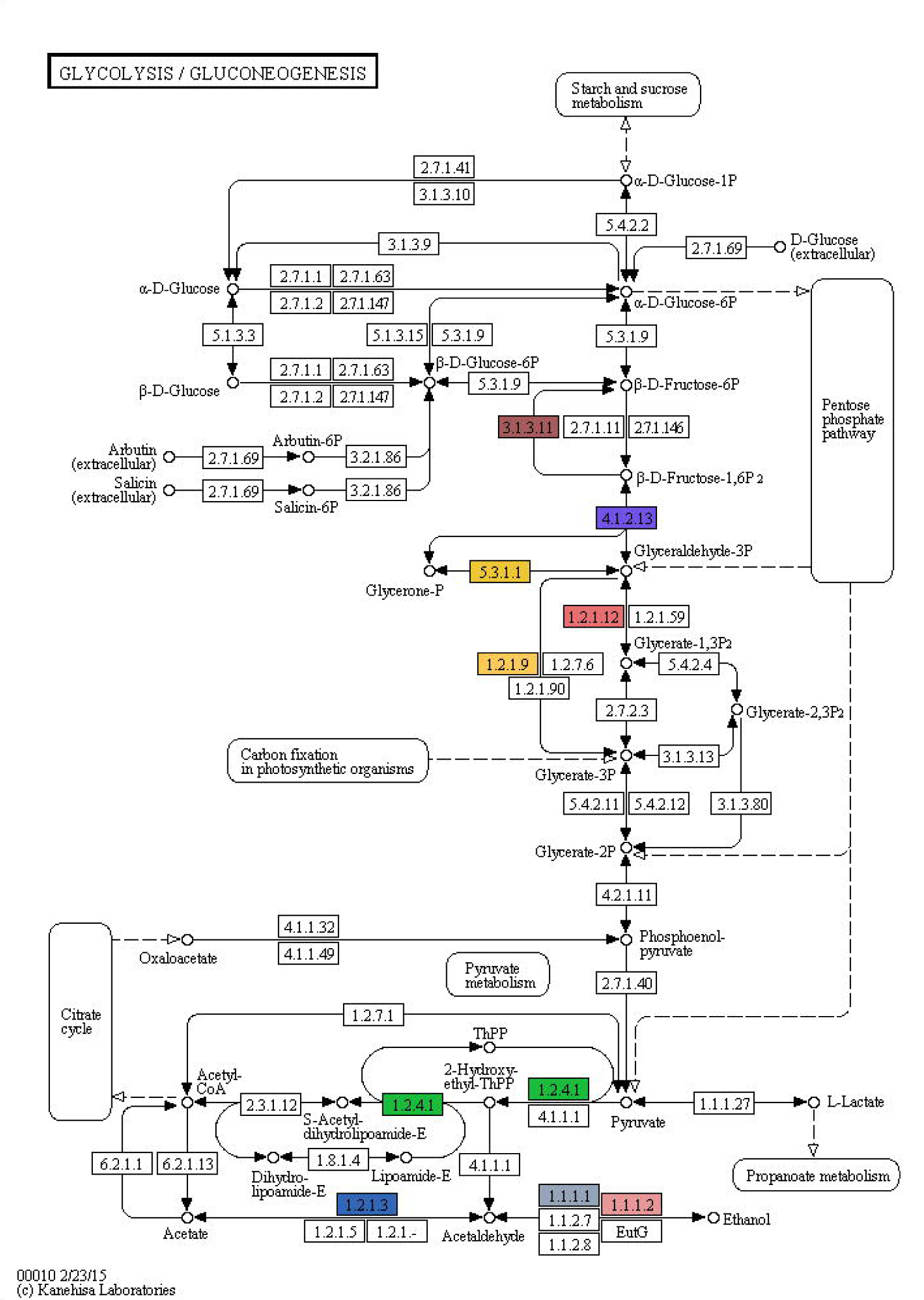

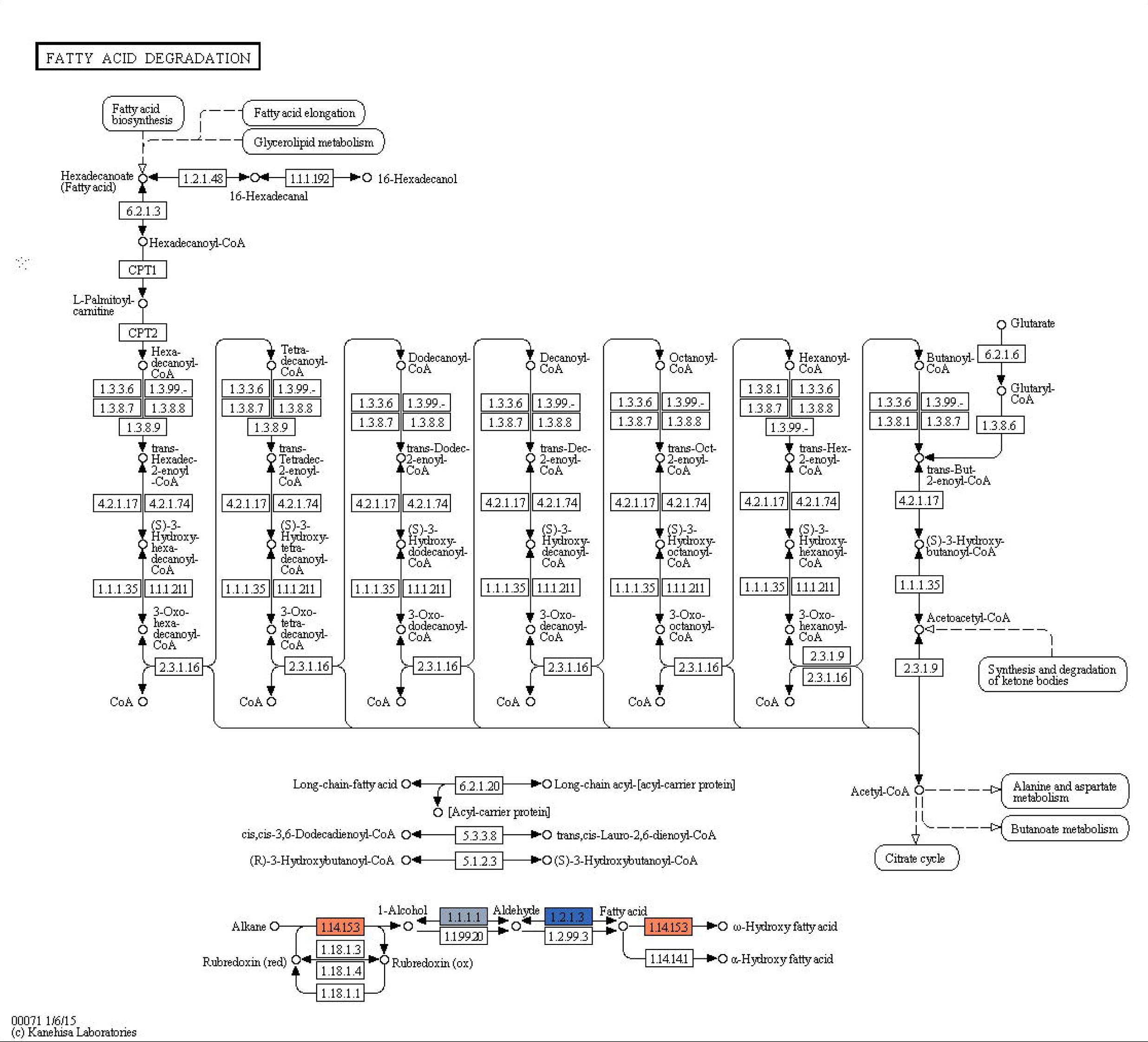

### Olive EST-SSR primers database

All primers were listed in the (**Table S3**) and was provided with all related information such as primer name, NCBI GI number for the EST sequence which is targeted by this primer, repeat type, repeat sequence, repeat length, repeat start index in the sequence, repeat end index in the sequence, forward and reveres primer pairs, annealing temperature (Tm) (°C), primer length (bp), primer product length (bp), the sequence of the EST, sequence description, gene ontology, enzyme code and enzyme name.

### Validation of designed primers

Twenty five primers were randomly selected to validate its efficacy to be used in polymerase chain reaction (PCR) procedures as a reliable molecular marker for marker-assisted selection programs by using a genomic DNA isolated from nine olive cultivars. All tested primers, exhibited successfully amplified and detectable PCR bands and only 10 exhibited detectable polymorphism (Figure 8).

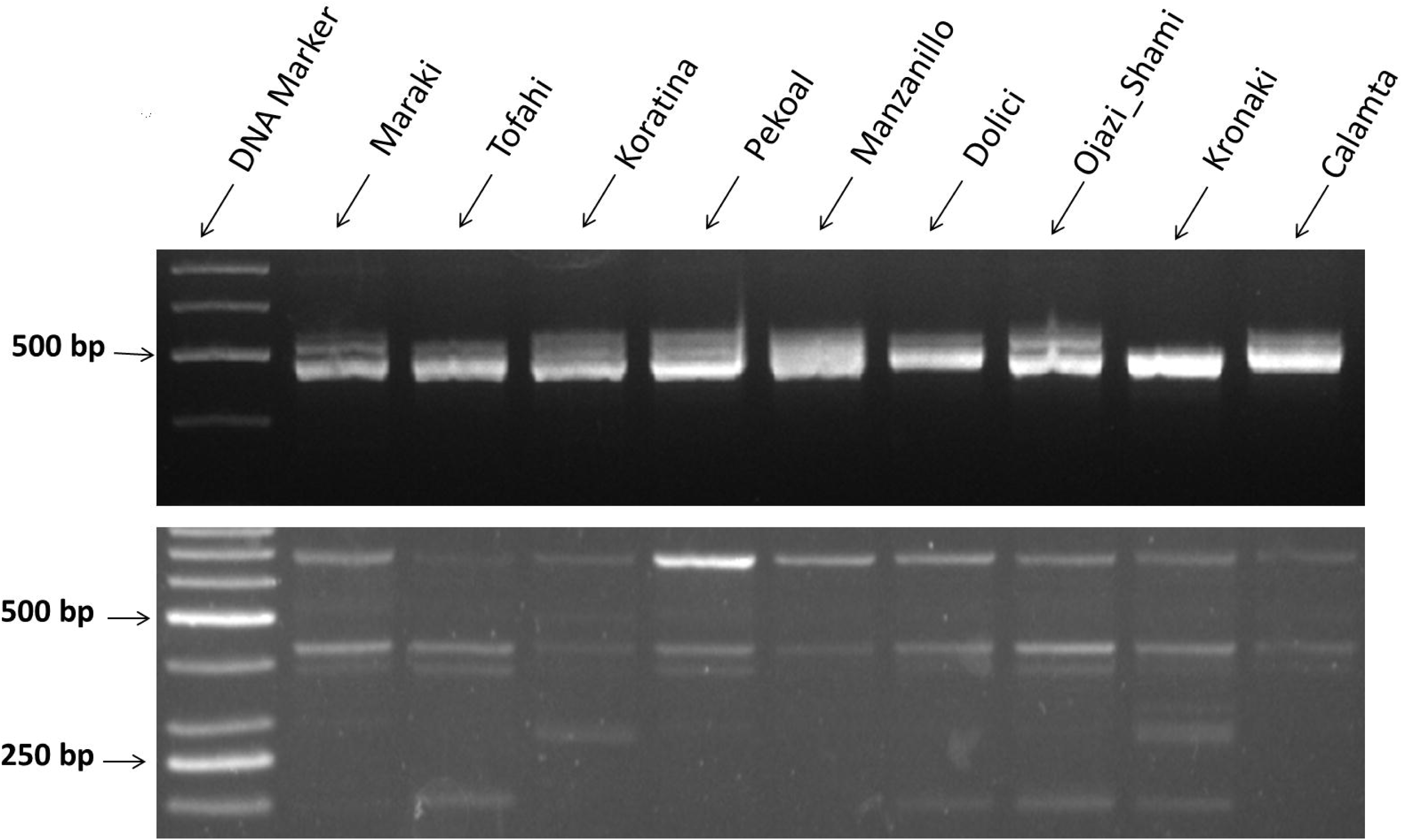

## Conclusion

SSR markers are very important co-dominant, highly polymorphic technique, which can be generated from functional regions in different plant genomes. The EST SSR technique has the potential to generate prototypically linked functional markers and it is a useful tool could be used in genetic diversity, marker assisted selection and genome mapping in olives. This study exhibits the functional categorization of olive EST sequences containing SSR motifs which can be targeted by a valid set of PCR primers. These ESTs representing genes associate with cellular component, biological process and molecular functions in olives. Also EST-SSR primers could provide useful information to understand the biological functions and gene-gene interactions by taking in advance the localization of these primers in different pathways which has possible relationships with highly important pathways in olive cultivation.

